# Simulated Plant Images Improve Maize Leaf Counting Accuracy

**DOI:** 10.1101/706994

**Authors:** Chenyong Miao, Thomas P. Hoban, Alejandro Pages, Zheng Xu, Eric Rodene, Jordan Ubbens, Ian Stavness, Jinliang Yang, James C. Schnable

## Abstract

Automatically scoring plant traits using a combination of imaging and deep learning holds promise to accelerate data collection, scientific inquiry, and breeding progress. However, applications of this approach are currently held back by the availability of large and suitably annotated training datasets. Early training datasets targeted arabidopsis or tobacco. The morphology of these plants quite different from that of grass species like maize. Two sets of maize training data, one real-world and one synthetic were generated and annotated for late vegetative stage maize plants using leaf count as a model trait. Convolutional neural networks (CNNs) trained on entirely synthetic data provided predictive power for scoring leaf number in real-world images. This power was less than CNNs trained with equal numbers of real-world images, however, in some cases CNNs trained with larger numbers of synthetic images outperformed CNNs trained with smaller numbers of real-world images. When real-world training images were scarce, augmenting real-world training data with synthetic data provided improved prediction accuracy. Quantifying leaf number over time can provide insight into plant growth rates and stress responses, and can help to parameterize crop growth models. The approaches and annotated training data described here may help future efforts to develop accurate leaf counting algorithms for maize.

## Introduction

Collection of trait data has become the rate-limiting step in many aspects of quantitative genetics of plant and animal breeding [1]. high-throughput phenotyping technologies hold promise to address this bottleneck [2–4]. The various technologies grouped under this header seek to either increase throughput of collecting trait data and/or decrease the cost per data point collected. One commonly stated goal is to do for trait data what advances in high-throughput sequencing technology did for genetic data. Yet, advances in sequencing technology alone would not have changed biology. New software tools, statistical methods, and experimental approaches were also required. The same is true for high-throughput phenotyping. It will not be enough to build the technology only to collect large amounts of raw data. New software methods are also needed to convert raw output from RGB, hyperspectral, LIDAR or other sensors into quantitative or qualitative measurements of plant traits.

Approaches to extracting quantitative or qualitative measurements of plant traits from images or other sensor data (such as point clouds produced by LIDAR sensors) can be broadly divided into two categories: 1) those where the programmer or scientist is responsible for telling the computer how to process the images [5] and 2) those where the computer learns how to process images from sets of annotated training data [6, 7]. Both approaches require a combination of image/sensor data and ground truth measurements. The first set of approaches may require greater investments of skilled labor in designing and troubleshooting algorithms, but smaller sets of ground truth measurements to obtain suitable accuracy of measurement [5, 8–11]. The second set of approaches, based on deep machine learning, often require substantially larger ground truth datasets to act as training data, but smaller investments of programmer time per additional trait scored [12–16]. The use of training datasets which were either insufficient in size or those with low quality annotations can lead to low prediction accuracy and “over fitting”, where training results in excellent prediction accuracy on the training data itself but low prediction when applied to newly collected data.

A number of approaches have been evaluated for generating the large and well annotated training datasets necessary for training deep learning algorithms to score traits from the raw datasets collected by different high-throughput phenotyping technologies. One such approach is to utilize crowdsourcing to outsource the generation of manual human annotations at a low cost per image. Paid crowdsourcing through Amazon’s Mechanical Turk platform has been used to identify and annotate maize tassels [17]. Observers working through the Zooniverse platform can achieve accuracy in arabidopsis leaf identification equivalent to that achieved by expert annotators [18]. Recently, a second approach to the cost-effective generation of large datasets has begun to attract interest in the plant phenotyping world: the use of computer generated synthetic training datasets. Simulation can produce large numbers of annotated training images at a small fraction of the cost of conventional data generation. As the three-dimensional structure and properties of the simulated plants are predetermined, accurate annotation of numerous traits is possible without time intensive and costly human intervention. A deep learning approach trained entirely on simulated data was shown to accurately count fruits in images collected from real-world plants [19]. The use of simulated training data has also been shown to increase the accuracy of both leaf counting in arabidopsis [16, 20] and the estimation of characteristics of three-dimensional root systems from two-dimensional images [21]. Outside of plant science, simulated training data has been used to train neural nets to recognize different stages of galaxy development in astronomical datasets [22]. The use of simulated image data ameliorates one potential bottleneck: the generation and annotation of training datasets. However, it often creates another: the creation of accurate *in silico* models of different plant species, which can replicate naturally occurring variation in target traits with sufficient fidelity to provide accurate training data for machine learning algorithms.

In this study we evaluate the feasibility of adapting existing procedural plant structural models, originally developed for creating scenery and backgrounds in video games and other computer graphics applications [23], to generate training data for machine learning algorithms intended to analyze real-world high-throughput plant phenotyping data. We demonstrate that simulated data can be used to generate models to estimate leaf number in real-world images, although training with a given number of simulated images provides lower accuracy than training with an equivalent number of accurately annotated real-world images. We further demonstrate that when annotated real-world images are scarce, increasing the size of training datasets by adding simulated data can produce a significant increase in prediction accuracy. Our results suggest that using simulated images as the training data may become an important tool for increasing the number of plant traits which can be effectively scored from images, particularly when well annotated real-world training data is absent or scarce.

## Results

An initial set of real-world maize plant images was generated using the University of Nebraska-Lincoln Greenhouse Innovation Center’s (UNL-GIC) automated greenhouse facility [24, 25]. A single replicate of the Buckler-Goodman maize association panel, a population selected to capture as much of genetic diversity segregating in cultivated maize as possible, was grown at the UNL-GIC [26]. Here we employ RGB images captured from 281 unique plants at 2-3 day intervals between 20 and 66 days after planting. A set of 6,620 RGB images taken from an angle approximately perpendicular to the axis on which most leaves emerged was uploaded to the Zooniverse crowdsourcing platform. Users were directed to exclude images presenting major challenges to accurate leaf counting, resulting in 4,633 RGB maize images with associated leaf count annotations [27]. Photos of early-stage plants taken at a higher zoom level were excluded from downstream analyses (1,212 images), as were 171 images where the number of observed leaves was <8 or >16. Leaf counts outside of the range of 8-16 were observed only infrequently (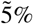 of cases). Thus the final dataset consists of a total of 3,250 annotated images from genetically diverse maize plants in the adult vegetative stages or early reproductive stages of development.

The experimental design for annotation included a modest level of repeated image scoring by the same user, and a more significant level of repeated scoring of the same image by multiple users. These overlaps enabled tests of both consistency in scoring by individuals, and agreement in scoring across individuals. Leaf counts applied to the same image presented to the same worker at different times were highly consistent (*r*^2^ = 0.94) (Figure 1A). Agreement of leaf counts between different workers scoring the same images was somewhat lower (*r*^2^ = 0.87) (Figure 1B), and, notably, individual workers exhibited bias relative to each other, with some tending to produce higher leaf counts than others from the same images (Figure S1).

**Figure 1.**
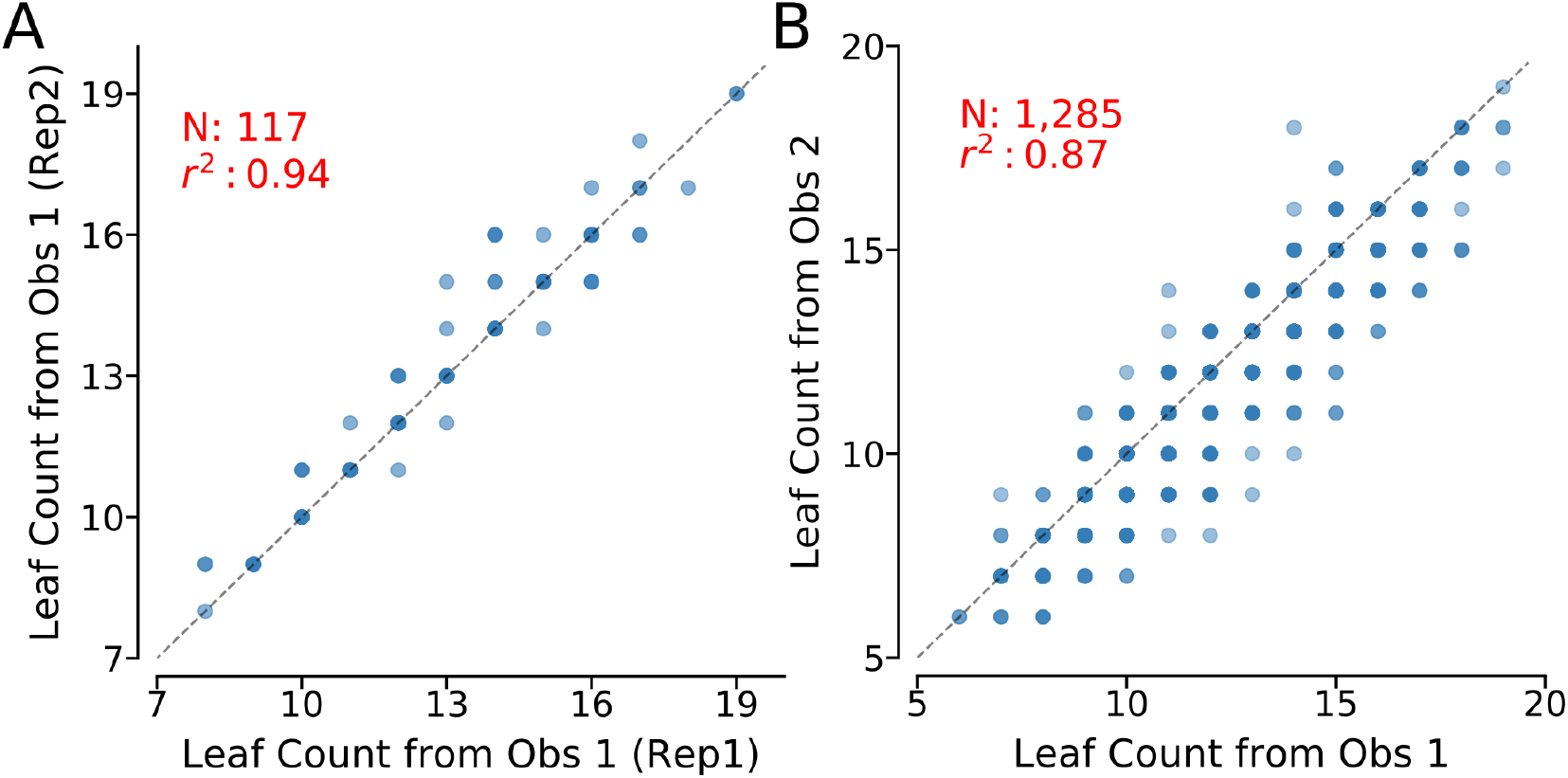
Intra- and inter-observer variation in leaf counting results. (A) Agreement in maize leaf counting results for the same image shown to the same observer at different times. (B) Agreement in maize leaf counting results for the same image shown to two different observers. Darker circles indicate a higher density of datapoints with the same x-axis and y-axis values. Squared correlation coefficients (*r*^2^) and the number of datapoints are reported for both panels. Obs: Observer.

Manual follow-ups were able to trace many disagreements between observers to one of four issues:

1. Differences in when a new leaf, emerging from the top of the whorl, was included in the leaf count for the plant (Figure 2A).
2. Differences in whether senescing, juvenile and damaged leaves at the base of the stalk were still included in the leaf count for the plant (Figure 2B).
3. Leaves extending either directly away from or toward the perspective of the camera, causing the leaf to either occlude or be occluded by the stalk (Figure 2C).
4. Partially or completely overlapping leaves from the perspective of the observer (Figure 2D).

**Figure 2.**
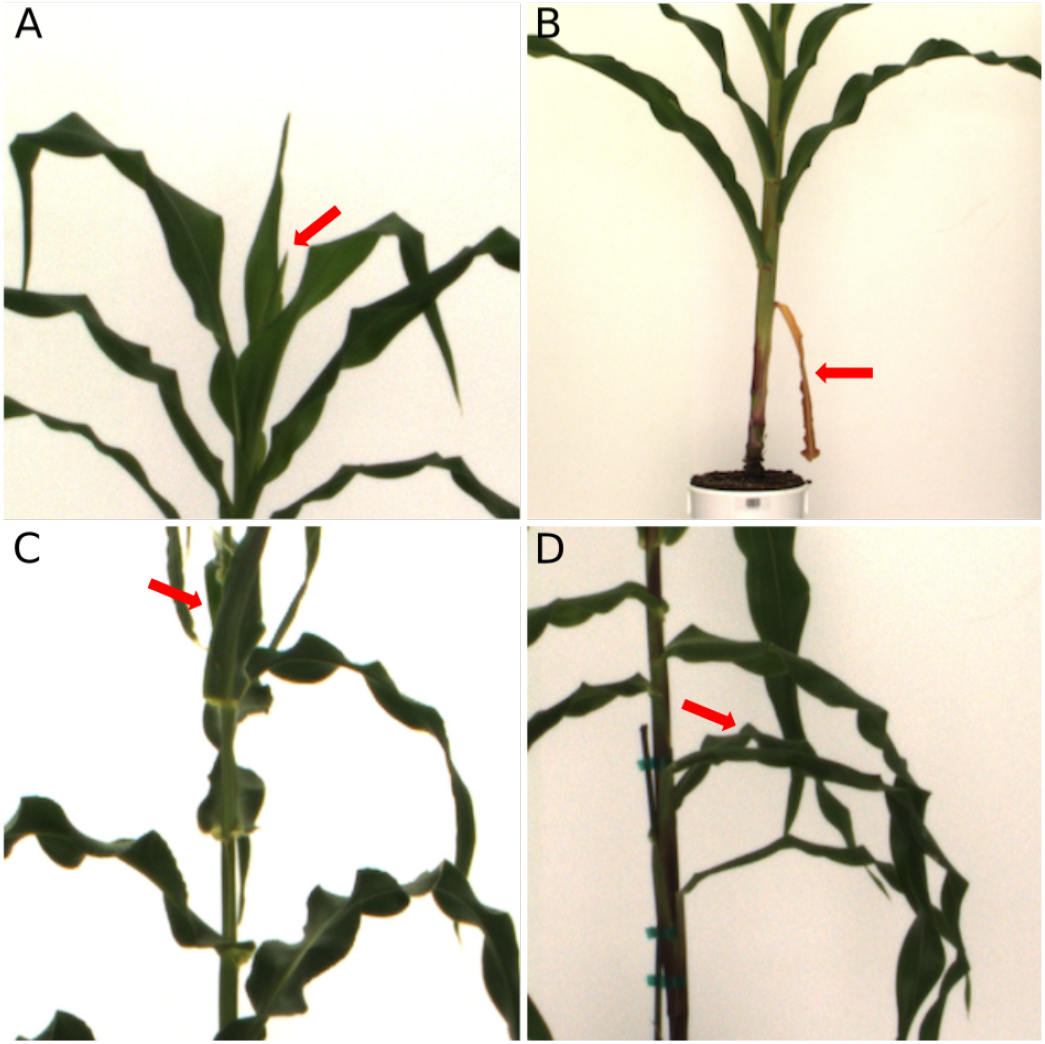
Common sources of disagreements in reported leaf number for the same image. In each panel the area of the problematic feature is indicated with a red arrow. A) The tip of a new leaf just beginning to emerge from the central whorl. B) A damaged and senescing leaf at the base of the stalk. C) Leaves emerging parallel with the angle of observation rather than perpendicular to it. D) Leaves which partially or completely overlap in this particular camera angle.

The first and second of these issues can be addressed in future labeling experiments for maize by adopting more explicit instructions now that common points of confusion have been identified. The third and fourth of these issues are unavoidable challenges of attempting to quantify three-dimensional structures, such as plant architectures, using two-dimensional data from a single perspective.

Using this initial manually annotated training data, a range of models were trained using sub-sampled training datasets with equal representation of each leaf count category between 8 and 16 leaves, i.e. 45-720 training images. Model accuracy, quantified both based on *r*^2^ and RMSE improved as the size of the training data increased (Figure 5A and C). Using the complete training data set of 720 images, prediction accuracy was *r*^2^ = 0.74 and RMSE = 1.33 (Figure 3) on separate test data.

**Figure 3.**
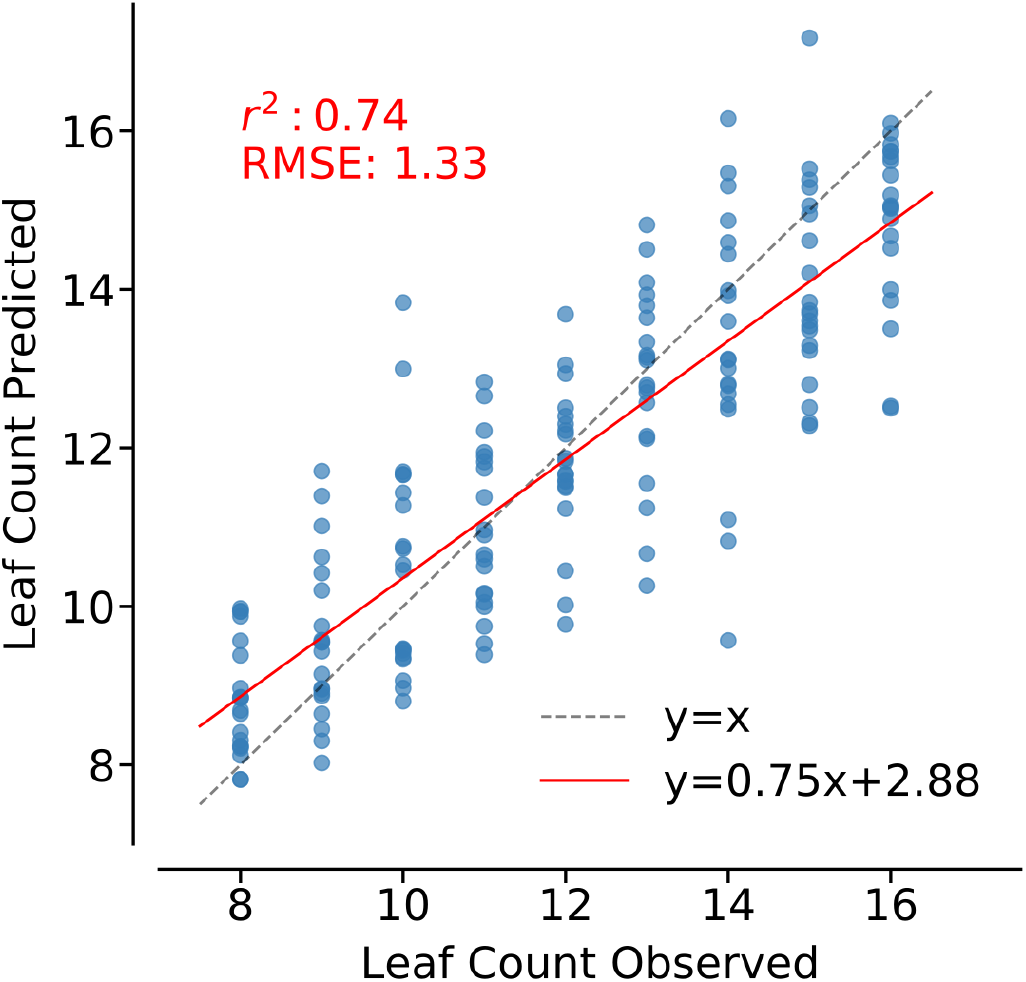
Prediction accuracy of a model trained using 720 real-world maize images. Correlation between observed and predicted leaf number for a separate set of 180 images (20 per leaf number category). The performance of the model (*r*^2^ and RMSE) was indicated at the upper left corner. Perfect prediction accuracy – a one-to-one relationship between observed and predicted values – is shown as a dashed gray line. The best-fit linear regression line for this data and the equation for this line are indicated in red.

In parallel with manual scoring of real-world maize images, a second dataset was procedurally generated using simulated maize plant architectures created using the maize module for Plant Factory Exporter [23] (see Methods). The resulting synthetic images were comparable to real-world images in a number of ways, although not to a degree that would lead to any risk of confusion by a human observer (Figure 4). Maize plants tend to have imperfect and alternating phylotaxy with successive leaves tending to emerge on opposite sides of the stalk. This pattern of leaf placement was not accurately captured in the commercially available procedural model where successive leaves emerged from the stalk at random angles (Figure 4). Synthetic data with a wider range of leaf numbers were initially generated, however a subset of 3,655 images with at least two hundred images of simulated plants in each leaf number between 8 leaves and 16 leaves was employed for downstream analyses.

**Figure 4.**
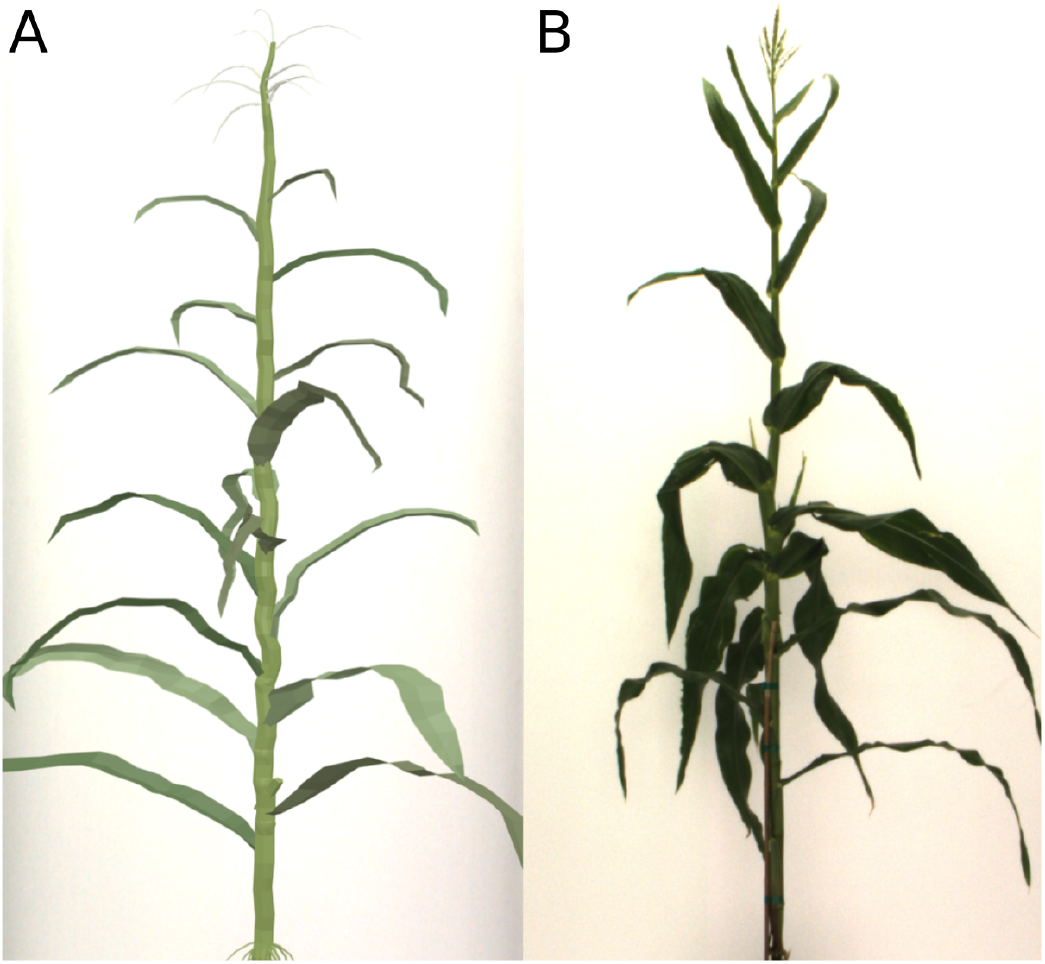
Comparison of synthetic and biologically derived maize plant images. (A) An example of a simulated photo of a maize plant generated using Blender and Plant Factory Exporter for this study (see Methods). (B) An example of a real-world photo of a maize plant used in this study.

Models were trained as described above, but using entirely synthetic data and these models were evaluated for their accuracy in determining leaf numbers from real-world maize images. Models trained on purely synthetic data provided lower prediction accuracy than models trained on real-world data given the same amount of training data (Figure 5A, S2). However, in some cases the accuracy of models trained using larger synthetic dataset did outperform the accuracy of models trained using smaller real-world datasets (Figure 5A, S2). The latter comparison may have more real-world relevance as the monetary, labor, and time costs of generating even a small real-world dataset with high quality annotations dramatically exceed those of generating even a large synthetic dataset.

**Figure 5.**
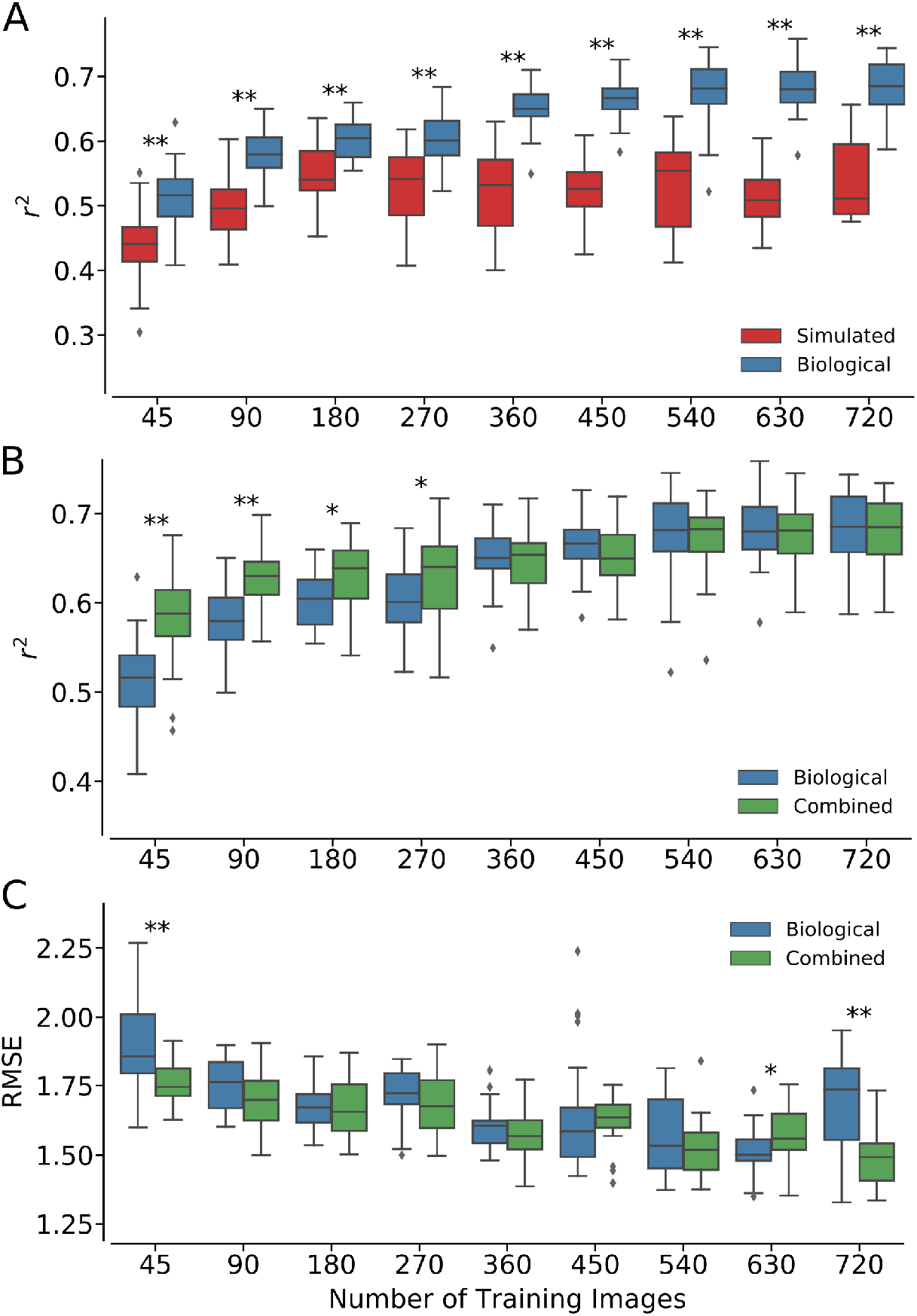
Performance of leaf counting models trained using synthetic, biologically derived, or combined training data. (A) Change in *r*^2^ prediction accuracy in response to increased training data size for models trained using either purely simulated or purely biologically-derived maize images. (B) Change in *r*^2^ prediction accuracy in response to increased training data size for models trained using either purely biologically-derived maize images or combined simulated and biological training data. (C) Change in RMSE prediction accuracy in response to increased training data size for models trained using either purely biologically derived maize images or combined simulated and biological training data. In all three panels, statistically significant differences are indicated using either * (*p* values < 0.05) or ** (*p* values < 0.01) (two-sided t-test).

We next evaluated whether using synthetic training data to augment real-world training data provides improvements in prediction accuracy. Two hundred simulated images per leaf number were added to the subsampled real-world training data. Models trained on the combined dataset exceeded the *r*^2^ accuracy of models trained on purely real-world data when real-world training data was rare (≤ 270 real-world images). When real-world training data was more abundant, models trained on combined datasets provided equivalent *r*^2^ accuracy to models trained on purely real-world data when real-world training data was more abundant (Figure 5B). Even after the *r*^2^ accuracy of models trained with real-world and combined real-world and synthetic training data converged, models trained using combined data included to show advantages in RMSE in some cases (Figure 5C). This suggests that the addition of synthetic data may reduce over/under prediction, as has been previously reported for synthetic *A. thaliana* training datasets [16].

## Discussion

Successfully moving the field of plant phenotyping forward requires bringing together engineers, computer scientists, and plant biologists to work on common problems. Interest in the development and improvement of computer vision approaches to leaf counting has been one of the early successes in recruiting computer scientists to work on a computational problem within the sphere of plant science and agriculture. Initial work in this field has focused on rosette plants, specifically arabidopsis and tobacco. While arabidopsis is a widely used genetic model, a great deal of the credit for this initial focus must go to the creators of the original annotated aArabidopsis and tobacco dataset [28] as well as the organizers of the Leaf Segmentation Challenge as part of the recurring Computer Vision Problems in Plant Phenotyping workshop [29]. These early successes provide a partial roadmap for how plant scientists can work to make other computer vision problems in plant science and agriculture appealing to computer scientists. Here we have generated a set of 3,421 annotated images of maize plants, capturing a large proportion of the genetic and phenotypic diversity present within temperate maize germplasm [26].

Given the similarity of leaf and plant architectures of arabidopsis and tobacco, some algorithms have been shown to provide useful results from cross-training and/or training on combined datasets [30]. arabidopsis and tobacco share rosette-style leaf architectures prior to reproductive transition, enabling effective capture of the majority of non-occluded leaves in a single top-down photo [28]. The leaf morphologies of both species are also similar, with a narrow petiole which then expands into the leaf blade. Maize belongs to the grass family (Poaceae), one of the most ecologically successful and agriculturally important clades of plants on the planet today. Almost fifty different grasses have been domesticated, thirty as sources of grain [31]. Of these grain crops, three of those grain, three (maize, rice, and wheat) are responsible, for half of all calories that sustain the human population around the world in an average year, either through direct consumption or indirectly after being fed to livestock. The grasses exhibit a radically different plant architecture from plants such as arabidopsis and tobacco. Leaves emerge from nodes along an elongated stalk and are separated into the leaf sheath, generally wrapped around the stalk, and the leaf blade, the portion of the leaf which extends away from the stalk. The widest portion of the leaf will generally occur right where the leaf meets the stalk. The base of each leaf sheath is often obscured beneath the sheath of leaf emerging from the previous node. In many cases, grasses also produce tillers – multiple independent stalks – each bearing a separate set of leaves. When leaf counting is possible, it is generally attempted from a side perspective, as was done here, and not from the top down, as is often done for arabidopsis. Advances in automated leaf counting of the grasses, including maize, can directly translate into beneficial outcomes for breeding and agronomic prediction Leaf initiation, leaf appearance rate, and final leaf number all vary between environments, between genotypes, and exhibit nonadditive genotype by environment interactions [32]. Parameterizing genotype-specific crop growth models requires knowledge of these parameters [33–35]. A decrease in the rate of new leaf emergence can provide an early warning sign of plant stress, and its detection can aid in variety selection in a breeding context, and guide early interventions to protect crop health and yield in a precision agriculture context.

More training data will, almost, always improve prediction accuracy for any computer vision task. However, the generation and labeling of training data can often become rate-limiting. This is particularly true for larger crop plants, such as maize and sorghum, where the logistics of growing and imaging plants in a controlled setting rapidly become challenging as the number of individuals being grown and imaged increases. As discussed above, simulated data has been shown to aid in improving the accuracy and reducing the biases of models trained to score plant traits from images [16, 19–21]. However, the creation of accurate plant morphological models to create these simulated datasets can be challenging. Here we evaluated the utility of simulated images from a commercially available procedural model. Despite numerous biological inaccuracies (Figure 4), training using purely simulated images provided non-zero accuracy, demonstrating that simulated images can act as one potential substitute when real-world training data is entirely unavailable, particularly when the simulated dataset can capture a superset of the geometries expected to be observed in real-world plants rather than a subset. More encouragingly, we found that using simulated images to augment real-world training data provided an increase in accuracy, particularly when real-world training data was rare (Figure 5B and C). In this example, we found that the benefit of synthetic data augmentation decreased as the size of the real-world training dataset increased.

Going forward, there are four clear avenues to further improve prediction accuracy. The first is to improve the accuracy of synthetic plant images. The commercially available procedural model employed here was sufficient for proof of concept, however there is a great deal of scope for further improvements in biological accuracy including the corporation of gene networks and weather models [36], as well as simulating how plants grow and interact with their environments over time [37, 38] to produce final adult structures more comparable to those observed in real-world plants. The second is to experiment with a wider range of potential models. For simplicity of comparison, here we adopted a previously proposed CNN model for plant phenomics including a total of four convolutional layers [14]. However, models with more layers and/or more complex model structures could also be evaluated, as these have been shown to provided higher prediction accuracy in some use cases [7, 30]. The third limitation in the present study is the quality of available training data (Figure 1). Current image labels reflect a modest degree of observer bias S1 and even perfect prediction accuracy will appear imperfect in the presence of errors in ground truth scoring. Labeling of each image by multiple observes would enable correction for this bias. However, some errors in true leaf number of unavoidable in either human or computational scoring of two-dimensional images as some leaves will be impossible to observe from any given perspective (Figure 2). Automated greenhouse technology makes it straightforward to image the same plant from multiple angles. Achieving highly accurate leaf number scoring of maize – as well as related crops such as sorghum, rice, and wheat – will require integrating images of the same plant collected from multiple viewing angles, either being fed into a CNN model directly, or an intermediate stage with a three-dimensional model is reconstructed prior to leaf counting. The challenges of acquiring large numbers of accurate three-dimensional structures from real-world plants may make synthetic plant datasets even more useful as a source of training data in that domain.

## Methods

### Collecting real-world Maize Images

A set of 281 lines from the maize Buckler-Goodman association panel [26] where grown in the greenhouse of the University of Nebraska-Lincoln’s Greenhouse Innovation Center (Latitude: 40.83, Longitude: −96.69) [24, 25] between August 1st to October 11th, 2018 and imaged from August 17th to October 7th, 2018. Kernels were sown in 2.4 gallon pots with Fafard germination mix supplemented with 1 cup (236 mL) of Osmocote plus and 1 tablespoon (15 mL) of Micromax Micronutrients per 2.8 cubic feet (80 L) of soil. The target photoperiod was 14:10 with supplementary light provided by light-emitting diode (LED) growth lamps from 07:00 to 21:00 each day. The target temperature of the growth facility was between 24–26°C. After growing in the greenhouse for seventeen days, all the plants were moved on to the conveyor belt which transferred each pot to the imaging chamber every two days and to the watering station each day. At the watering station plants were weighed once per day and watered back to a target weight, including pot, soil, carrier, and plant of 7,500 grams on August 17th, 7,400 grams from August 18th to September 2nd, 7,300 grams from September 3rd to September 16th, 7,800 grams on September 17th, 8,300 grams from September 18th to September 27th, and 9,000 grams from September 28th to the termination of the experiment. This increase in target weight at later days reflects the increasing demand for water from the maturing maize plants.

Plants were imaged every 2-3 days between 29 and 66 days after planting using the set of high-throughput phenotyping imaging systems in the previously-described greenhouse facility [24, 25]. If possible plants were arranged so that most leaves were perpendicular to the line between the camera and the center of the stalk. A total 6,620 RGB images were captured at a resolution of 2,454 × 2,056 pixels. At the zoom level used in this paper, each pixel represented an area of approximately 2.36 mm^2^ for objects in the range between the camera and the pot containing the plant.

### Creating Synthetic Maize Images

Initial three-dimensional maize plant structures were generated using the “maize/corn” module within Plant Factory Exporter (Plant Factory Exporter 2016 R3) with each parameter of the procedural model assigned a new random value for each new three-dimensional model [23]. Parameters for the “maize/corn” module included plant height, leaf number and placement, leaf length, leaf angle, leaf smoothness, stem thickness, number of prop root whorls, number of tassel branches, etc. A three-dimensional recreation of the imaging chamber was constructed in Blender (v2.69) with an image of the chamber taken without a plant used to provide two-dimensional textures for the back wall and the plant carrier/pot. The three-dimensional structures created in Plant Factory Exporter were imported into Blender, resized and recolored based on the height and color range observed from real plants, and any ears placed by Plant Factory Exporter were deleted. three-dimensional maize plant models were rotated within the three-dimensional imaging chamber to allow simulated photography from multiple angles. The importing, resizing, recoloring, ear deletion, rotation and capture of two-dimensional images were automated using Blender’s Python scripting interface. This script is provided as part of the associated GitHub repository for this paper.

### Scoring Leaf Numbers

Real-world maize images were scored for leaf number using the Zooniverse crowdsourcing platform. The leaf counting project was created on Zooniverse website (https://www.zooniverse.org/projects/alejandropages/maize-leaf-count) and all real images were uploaded using custom python scripts provided at the GitHub repository associated with this paper (Figure S3). A tutorial on how to do the counting was also set up to guide people to finish the leaf counting tasks. Observers were invited to count leaf numbers by clicking anywhere within the extent of each leaf in each photo. Prior to counting, observers were first asked to classify each image as scorable or unscorable image. Unscorable images included those without a plant, plants so tall that the top of the plant was out of frame, and plants with one or more tillers in addition to the primary stem. Images classified as unscorable did not proceed to the leaf counting step. Data from a total of four observers were used in downstream analysis, with 4,641, 1,917, 445, and 171 images scored by the most prolific to the least prolific observers respectively.

### Training and Evaluation of CNN Models

Training of leaf counting models was conducted using the Deep Plant Phenomics (DPP) package, which provides a plant phenotyping specific interface and wrapper on top of TensorFlow (v1.10) [14, 39]. Prior to both model training and testing, the input size of each image was set to 256 × 256 pixels. For all models trained changes in brightness and contrast, mirroring of the image, and cropping were all employed to augment the size of the training datasets. The model structure contained four convolutional layers with filter dimension (5, 5, 3, 32), (5, 5, 32, 64), (3, 3, 64, 64) and (3, 3, 64, 64) respectively. The filter dimension is in the order [x_size, y_size, depth, num_filters] where x_size and y_size are spatial dimensions, depth is the full depth of the input volume which is 3 for RGB images, and num_filters is the desired number of filters in this layer. The stride length was set to 1 for all the convolutional filters. A max pooling layer with 3 × 3 kernel size and a stride value of 2 were added following each convolutional layer to decrease spatial resolution. All experiments employed a static learning rate of 0.0001 and an epoch number of 500.

A total of 9 scenarios representing various number of training datasets as shown in Figure 5 were evaluated in each training image types including real, synthetic and combined. For each scenario, five unique models were trained using different subsets of training images with the same number of images per leaf number category. Every time the real images were selected for training a model, five subsets of testing data were also created from the remaining the images. In all scenarios, including those trained with real-world, synthetic, or combined data were evaluated using only real-world testing data to determine model accuracy. Thus, the accuracy of each of five models was tested in each of five testing datasets, which means a total of 5*5=25 distinct predictions were made for each scenario. With these predictions, the square of Pearson’s r (*r*^2^) and the Root Mean Square Error (RMSE) between predicted leaf numbers and actual leaf numbers were estimated for making the box plots as shown in the result section. All training and prediction analysis for this paper was conducted on nodes of the Holland Computing Center computational cluster at the University of Nebraska-Lincoln equipped with either Nvidia K20, K40, or P100 GPUs. All the Python scripts employed to automate and conduct training and predictions are made available in the GitHub repository associated with this publication.

## Acknowledgements

This work was supported by a University of Nebraska Agricultural Research Division seed grant to JCS, a National Science Foundation Award (OIA-1557417) to JCS and JY, and a UCARE fellowship to AP. This project was completed utilizing the Holland Computing Center of the University of Nebraska, which receives support from the Nebraska Research Initiative. This publication uses data generated via the Zooniverse.org platform, development of which is funded through multiple sources, including a Global Impact Award from Google, and by a grant from the Alfred P. Sloan Foundation.

## Author Contribution Statement

T.H. and J.C.S. generated the maize plant three-dimensional models and two-dimensional images. J.Y. and E.R. generated the real maize plant images. C.M. and A.P scored the real plant images. C.M. and Z.X. conducted the analyses. J.U. and I.S. advised on the approaches employed for deep learning. C.M. and J.C.S. wrote the paper. All authors reviewed and approved the manuscript.

## Additional Information

The scripts and source code employed in this study have been deposited at https://github.com/freemao/MaizeLeafCounting. Images and annotations used in this study have been deposited with CyVerse [27].

The authors declare no competing interests.

## Supplementary materials

**Figure S1.**
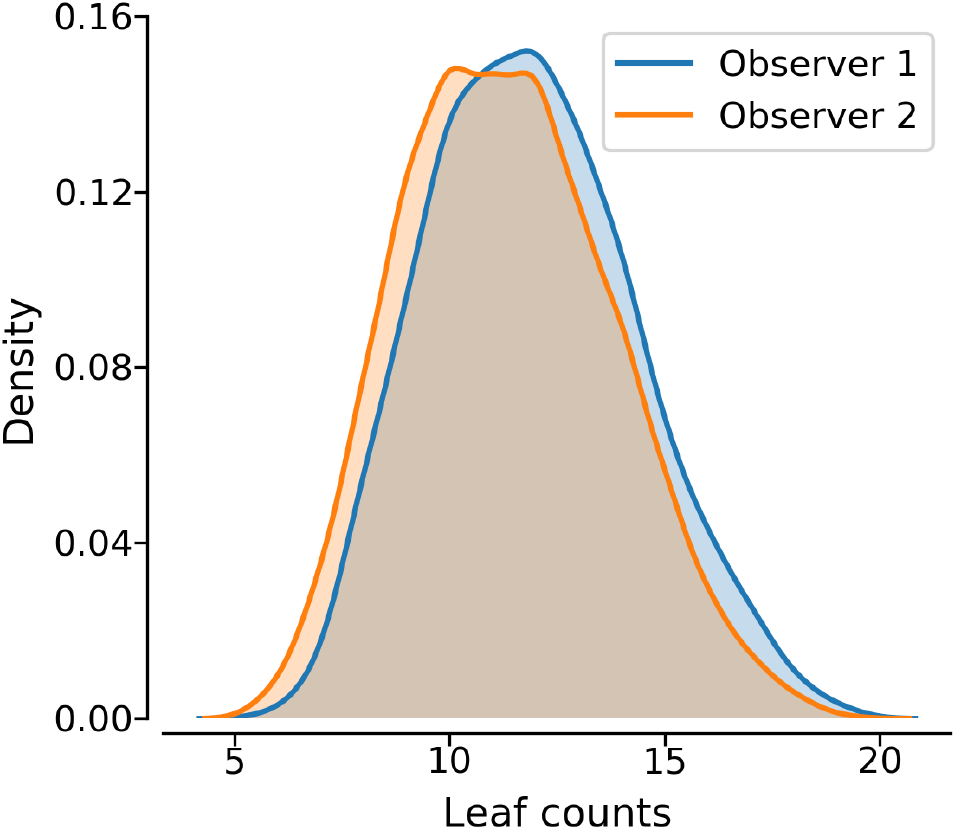
Differences is the distribution of reported leaf numbers from the two most prolific observers in this study.

**Figure S2.**
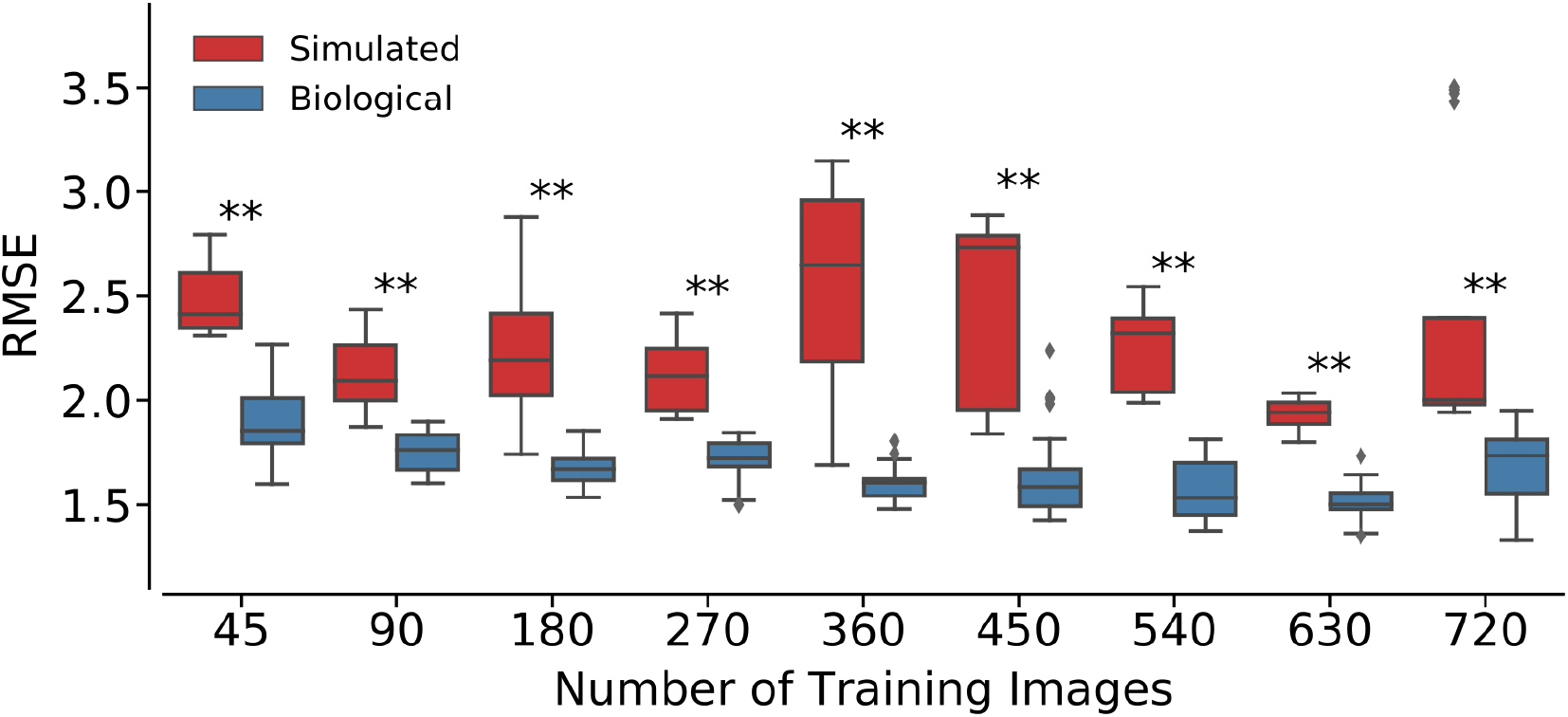
Changes in the RMSE of models trained using either purely real (biological) or purely simulated maize images across various number of images in the training dataset. The numbers on the x-axis represent the equivalent number of real or simulated images in each training dataset. Statistically significant differences are indicated with either * (p values < 0.05) or ** (p values < 0.01) (two sided t-test).

**Figure S3.**
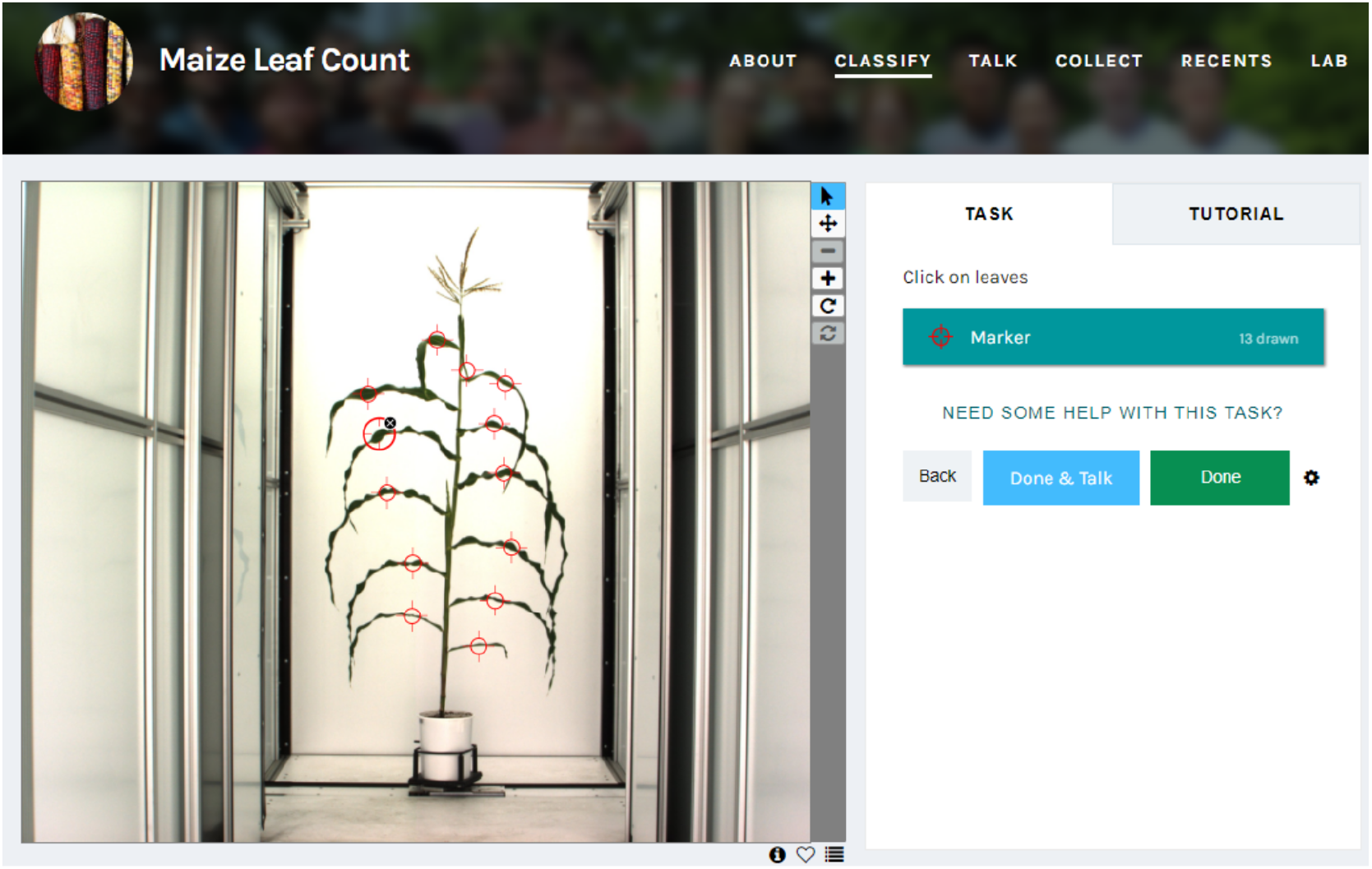
Maize leaf count Zooniverse project page. The text presented to each user when first starting the leaf counting project was: Tutorial Slide 1: “Welcome to Maize leaf count! Your task is to count the leaves of each plant. We have to filter out bad images first: Bad images are: Multiple plants in one pot, Serious Leaning, Too tall, Too unhealthy. (You can be conservative with the last one).” Tutorial Slides 2-5, examples of each kind of “bad” image. Tutorial Slide 6: “Leaf Count Use the leaf selection tool to count the leaves. Don’t worry about where to click, the tool is for you to keep track of which leaves you’ve already counted.” Tutorial Slide 7: “Thank you for your interest in our project! Please feel free to ask questions for more information.”

## References

1. Houle, D., Govindaraju, D. R. & Omholt, S. Phenomics: the next challenge. Nat. reviews genetics 11, 855 (2010).

2. Furbank, R. T. & Tester, M. Phenomics–technologies to relieve the phenotyping bottleneck. Trends plant science 16, 635–644 (2011).

3. Araus, J. L. & Cairns, J. E. Field high-throughput phenotyping: the new crop breeding frontier. Trends plant science 19, 52–61 (2014).

4. Fernandez, M. G. S., Bao, Y., Tang, L. & Schnable, P. S. A high-throughput, field-based phenotyping technology for tall biomass crops. Plant physiology 174, 2008–2022 (2017).

5. Gehan, M. A. et al. Plantcv v2: Image analysis software for high-throughput plant phenotyping. PeerJ 5, e4088 (2017).

6. Giuffrida, M. V., Minervini, M. & Tsaftaris, S. A. Learning to count leaves in rosette plants. (2016).

7. Aich, S. & Stavness, I. Leaf counting with deep convolutional and deconvolutional networks. In Proceedings of the IEEE International Conference on Computer Vision, 2080–2089 (2017).

8. Jansen, M. et al. Simultaneous phenotyping of leaf growth and chlorophyll fluorescence via growscreen fluoro allows detection of stress tolerance in arabidopsis thaliana and other rosette plants. Funct. Plant Biol. 36, 902–914 (2009).

9. De Vylder, J., Vandenbussche, F., Hu, Y., Philips, W. & Van Der Straeten, D. Rosette tracker: an open source image analysis tool for automatic quantification of genotype effects. Plant physiology 160, 1149–1159 (2012).

10. Fahlgren, N. et al. A versatile phenotyping system and analytics platform reveals diverse temporal responses to water availability in setaria. Mol. plant 8, 1520–1535 (2015).

11. Brichet, N. et al. A robot-assisted imaging pipeline for tracking the growths of maize ear and silks in a high-throughput phenotyping platform. Plant Methods 13, 96 (2017).

12. Mohanty, S. P., Hughes, D. P. & Salathé, M. Using deep learning for image-based plant disease detection. Front. plant science 7, 1419 (2016).

13. Pawara, P., Okafor, E., Surinta, O., Schomaker, L. & Wiering, M. Comparing local descriptors and bags of visual words to deep convolutional neural networks for plant recognition. In ICPRAM, 479–486 (2017).

14. Ubbens, J. R. & Stavness, I. Deep plant phenomics: a deep learning platform for complex plant phenotyping tasks. Front. plant science 8, 1190 (2017).

15. Pound, M. P. et al. Deep machine learning provides state-of-the-art performance in image-based plant phenotyping. Gigascience 6, gix083 (2017).

16. Ubbens, J., Cieslak, M., Prusinkiewicz, P. & Stavness, I. The use of plant models in deep learning: an application to leaf counting in rosette plants. Plant methods 14, 6 (2018).

17. Zhou, N. et al. Crowdsourcing image analysis for plant phenomics to generate ground truth data for machine learning. PLoS computational biology 14, e1006337 (2018).

18. Giuffrida, M. V., Chen, F., Scharr, H. & Tsaftaris, S. A. Citizen crowds and experts: observer variability in image-based plant phenotyping. Plant methods 14, 12 (2018).

19. Rahnemoonfar, M. & Sheppard, C. Deep count: fruit counting based on deep simulated learning. Sensors 17, 905 (2017).

20. Ward, D., Moghadam, P. & Hudson, N. Deep leaf segmentation using synthetic data. arXiv preprint arXiv:1807.10931 (2018).

21. Landl, M. et al. Measuring root system traits of wheat in 2d images to parameterize 3d root architecture models. Plant soil 1–21 (2018).

22. Primack, J. et al. Deep learning identifies high-z galaxies in a central blue nugget phase in a characteristic mass range. The Astrophys. J. 858, 114 (2018).

23. e-on Software. Plant factory exporter.

24. Ge, Y., Bai, G., Stoerger, V. & Schnable, J. C. Temporal dynamics of maize plant growth, water use, and leaf water content using automated high throughput rgb and hyperspectral imaging. Comput. Electron. Agric. 127, 625–632 (2016).

25. Liang, Z. et al. Conventional and hyperspectral time-series imaging of maize lines widely used in field trials. GigaScience 7, gix117 (2017).

26. Flint-Garcia, S. A. et al. Maize association population: a high-resolution platform for quantitative trait locus dissection. The Plant J. 44, 1054–1064 (2005).

27. Miao, C., Pages, A. & Schnable, J. C. Individual maize plant leaf count dataset. CyVerse DOI: https://www.doi.org/10.25739/f12d-tt60 (2019).

28. Minervini, M., Fischbach, A., Scharr, H. & Tsaftaris, S. A. Finely-grained annotated datasets for image-based plant phenotyping. Pattern recognition letters 81, 80–89 (2016).

29. Scharr, H. et al. Leaf segmentation in plant phenotyping: a collation study. Mach. vision applications 27, 585–606 (2016).

30. Dobrescu, A., Valerio Giuffrida, M. & Tsaftaris, S. A. Leveraging multiple datasets for deep leaf counting. In Proceedings of the IEEE International Conference on Computer Vision, 2072–2079 (2017).

31. Glémin, S. & Bataillon, T. A comparative view of the evolution of grasses under domestication. New phytologist 183, 273–290 (2009).

32. Padilla, J. & Otegui, M. Co-ordination between leaf initiation and leaf appearance in field-grown maize (zea mays): genotypic differences in response of rates to temperature. Annals Bot. 96, 997–1007 (2005).

33. Muchow, R. C., Sinclair, T. R. & Bennett, J. M. Temperature and solar radiation effects on potential maize yield across locations. Agron. journal 82, 338–343 (1990).

34. Technow, F., Messina, C. D., Totir, L. R. & Cooper, M. Integrating crop growth models with whole genome prediction through approximate bayesian computation. PloS one 10, e0130855 (2015).

35. Masjedi, A. et al. Sorghum biomass prediction using uav-based remote sensing data and crop model simulation. In IGARSS 2018-2018 IEEE International Geoscience and Remote Sensing Symposium, 7719–7722 (IEEE, 2018).

36. Marshall-Colon, A. et al. Crops in silico: generating virtual crops using an integrative and multi-scale modeling platform. Front. plant science 8, 786 (2017).

37. Stava, O. et al. Inverse procedural modelling of trees. In Computer Graphics Forum, vol. 33, 118–131 (Wiley Online Library, 2014).

38. Pirk, S., Niese, T., Hädrich, T., Benes, B. & Deussen, O. Windy trees: computing stress response for developmental tree models. ACM Transactions on Graph. (TOG) 33, 204 (2014).

39. Abadi, M. et al. TensorFlow: Large-scale machine learning on heterogeneous systems (2015). Software available from tensorflow.org.

